# Sequencing by avidity enables high accuracy with low reagent consumption

**DOI:** 10.1101/2022.11.03.514117

**Authors:** Sinan Arslan, Francisco J. Garcia, Minghao Guo, Matthew W. Kellinger, Semyon Kruglyak, Jake A. LeVieux, Adeline H. Mah, Haosen Wang, Junhua Zhao, Chunhong Zhou, Andrew Altomare, John Bailey, Matthew B. Byrne, Chiting Chang, Steve X. Chen, Claudia N. Dennler, Vivian T. Dien, Derek Fuller, Ryan Kelley, Omid Khandan, Michael G. Klein, Michael Kim, Bryan R. Lajoie, Bill Lin, Yu Liu, Tyler Lopez, Peter T. Mains, Andrew D. Price, Samantha R. Robertson, Hermes Taylor-Weiner, Ramreddy Tippana, Austin B. Tomaney, Su Zhang, Mark R. Ambroso, Rosita Bajari, Ava M. Bellizzi, Chris B. Benitez, Daniel R. Berard, Lorenzo Berti, Kelly N. Blease, Angela P. Blum, Andrew M. Boddicker, Leo Bondar, Chris Brown, Chris A. Bui, Juan Calleja-Aguirre, Kevin Cappa, Joshua Chan, Victor W. Chang, Katherine Charov, Xiyi Chen, Rodger M. Constandse, Ryan Costello, Weston Damron, Mariam Dawood, Nicole DeBuono, John D. Dimalanta, Laure Edoli, Keerthana Elango, Nikka Faustino, Chao Feng, Mathhew Ferrari, Keith Frankie, Adam Fries, Anne Galloway, Vlad Gavrila, Gregory J. Gemmen, James Ghadiali, Arash Ghorbani, Logan A. Goddard, Adriana R. Guetter, Garren L. Hendricks, Jendrik Hentschel, Daniel J. Honigfort, Yun-Ting Hsieh, Yu-Hsien Hwang Fu, Scott K. Im, Chaoyi Jin, Shradha Kabu, Daniel E. Kincade, Shawn Levy, Yu Li, Vincent K. Liang, William H. Light, Jonathan B. Lipsher, Tsung-li Liu, Grace Long, Rui Ma, John M. Mailloux, Kyle A. Mandla, Anyssa R. Martinez, Max Mass, Daniel T. McKean, Michael Meron, Celyne S. Moh, Rachel K. Moore, Juan Moreno, Jordan M. Neysmith, Cassandra S. Niman, Jesus M. Nunez, Micah T. Ojeda, Sara Espinosa Ortiz, Jenna Owens, Geoffrey Piland, Daniel J. Proctor, Josua B. Purba, Michael Ray, Daisong Rong, Virginia M. Saade, Sanchari Saha, Gustav Santo Tomas, Nicholas Scheidler, Luqmanal H. Sirajudeen, Samantha Snow, Gudrun Stengel, Ryan Stinson, Michael J. Stone, Keoni J. Sundseth, Eileen Thai, Connor J. Thompson, Marco Tjioe, Christy L. Trejo, Greg Trieger, Diane Ni Truong, Ben Tse, Benjamin Voiles, Henry Vuong, Jennifer C. Wong, Chiung-Ting Wu, Hua Yu, Yingxian Yu, Ming Yu, Xi Zhang, Da Zhao, Genhua Zheng, Molly He, Michael Previte

## Abstract

We present avidity sequencing - a novel sequencing chemistry that separately optimizes the process of stepping along a DNA template and the process of identifying each nucleotide within the template. Nucleotide identification uses multivalent nucleotide ligands on dye-labeled cores to form polymerase-polymer nucleotide complexes bound to clonal copies of DNA targets. These polymer-nucleotide substrates, termed avidites, decrease the required concentration of reporting nucleotides from micromolar to nanomolar, and yield negligible dissociation rates. We demonstrate the use of avidites as a key component of a sequencing technology that surpasses Q40 accuracy and enables a diversity of applications that include single cell RNA-seq and whole human genome sequencing. We also show the advantages of this technology in sequencing through long homopolymers.

## Main

Over the past 15 years, next generation sequencing (NGS) methods revolutionized a broad set of applications [1-8]. Multiple technologies have been introduced during this time, each having various strengths and limitations [9]. The technologies vary by accuracy, read length, run time, and cost. The most widely used method utilizes highly parallel and accurate short read sequencing described in Bentley et al. and termed sequencing by synthesis (SBS) [10].

The SBS methodology sequences DNA by controlled (i.e., one at a time) incorporation of modified nucleotides. [11]. The modifications consist of a 3’ blocking group and a dye label [12, 13]. The blocking group ensures that only a single nucleotide is incorporated, and the dye label enables each nucleotide to be identified following an imaging step. The blocking group and label are subsequently removed, completing the sequencing cycle. The cycle is repeated with the incorporation of the next blocked and labeled nucleotide. Incorporation of the modified nucleotide meets two objectives: to advance the polymerase along the DNA template and to differentially label the incorporated nucleotide for base identification. Although combining the two processes is efficient, it prevents the processes from being independently optimized. For example, further quality improvements could be achieved if different polymerases were used for stepping along the template and introducing a label. Also, costs could be reduced if reagent concentrations required to meet each objective could be varied independently. Principally, reagent costs increase because high yielding and rapid incorporation requires micromolar concentrations of the polymerase and nucleotides to drive the reaction [14-18]. The alternative of allowing longer incorporation times results in longer cycle times that have an additive effect over 300 cycles of step wise sequencing.

We present a novel sequencing chemistry, termed avidity sequencing, that separates and independently optimizes the controlled incorporation step and the nucleotide identification step to achieve increased base calling accuracy relative to SBS while reducing the concentration of key reagents to nanomolar scale. To advance this approach, we first had to overcome the technical challenge of signal persistence. For example, a potential strategy to separating the steps described above could be to first incorporate a 3’ blocked but unlabeled nucleotide and then to bind a complementary labeled nucleotide to the subsequent base in the template for base identification. This approach is problematic because the dissociation rate for single nucleotides from a polymerase-template complex is high, and the polymerase-nucleotide complex does not remain stable through imaging unless prohibitively high concentrations of nucleotides are present in the bulk solution. To overcome this challenge, we leveraged avidity.

Avidity refers to the accumulated strength of multiple affinities of individual non-covalent binding interactions, which can be achieved when multivalent ligands tethered in close proximity can simultaneously bind to their targets [19]. Coincident binding increases ligand affinity and residence time [20]. As an example of the dramatic impact avidity can have on both affinity and decreased dissociation rate, Zhang et al. demonstrated that by changing a monomeric nanobody to a pentameric nanobody, it is possible to achieve affinity gains and decrease dissociation rates by 3-4 orders of magnitude [21]. Our approach was to leverage avidity for nucleotide detection within the sequencing chemistry. We demonstrate here that avidity sequencing achieves accuracy surpassing Q40 and enables a diversity of applications that include single cell RNA-seq and whole human genome sequencing. We also demonstrate an improved ability of this chemistry to sequence through homopolymer sequences.

## Results

Prior to sequencing, DNA fragments of interest were circularized and captured on the surface of a flowcell. Clonal copies of the DNA fragments were then created through rolling circle amplification (RCA) generating approximately 1 billion concatemers on the flowcell surface [22-25]. The resulting concatemers, referred to as polonies using the original term coined by George Church [26], were used as the DNA substrate for sequencing. We then constructed the avidite: a dye labeled polymer with multiple, identical nucleotide attached. In the presence of a polymerase, the avidite was able to bind multiple nucleotides specifically in copies of a DNA fragment within a polony. A polymerase and a mixture of four avidites, each corresponding to a particular label and nucleotide, were applied to the flowcell and used for base discrimination. The avidite was not incorporated, but provided a stable complex, while enabling removal under specifically designed wash conditions. Removal of the avidite left no modifications in the synthesized strand. The avidites decreased the required concentration of reporting nucleotides by 100x relative to single nucleotide binding, yielded negligible dissociation rates, and obviated the need to have nucleotides present in the bulk solution. The advent of the avidite enabled us to separate the process of stepping along the DNA template from the process of identifying each nucleotide and to optimize each for quality and reagent consumption. Figure 1A shows a complete cycle of avidity sequencing. Figure 1B depicts a single avidite interacting with multiple DNA copies within a polony, and figure 1C shows many avidites specifically bound to several polonies on the surface.

**Figure 1:**
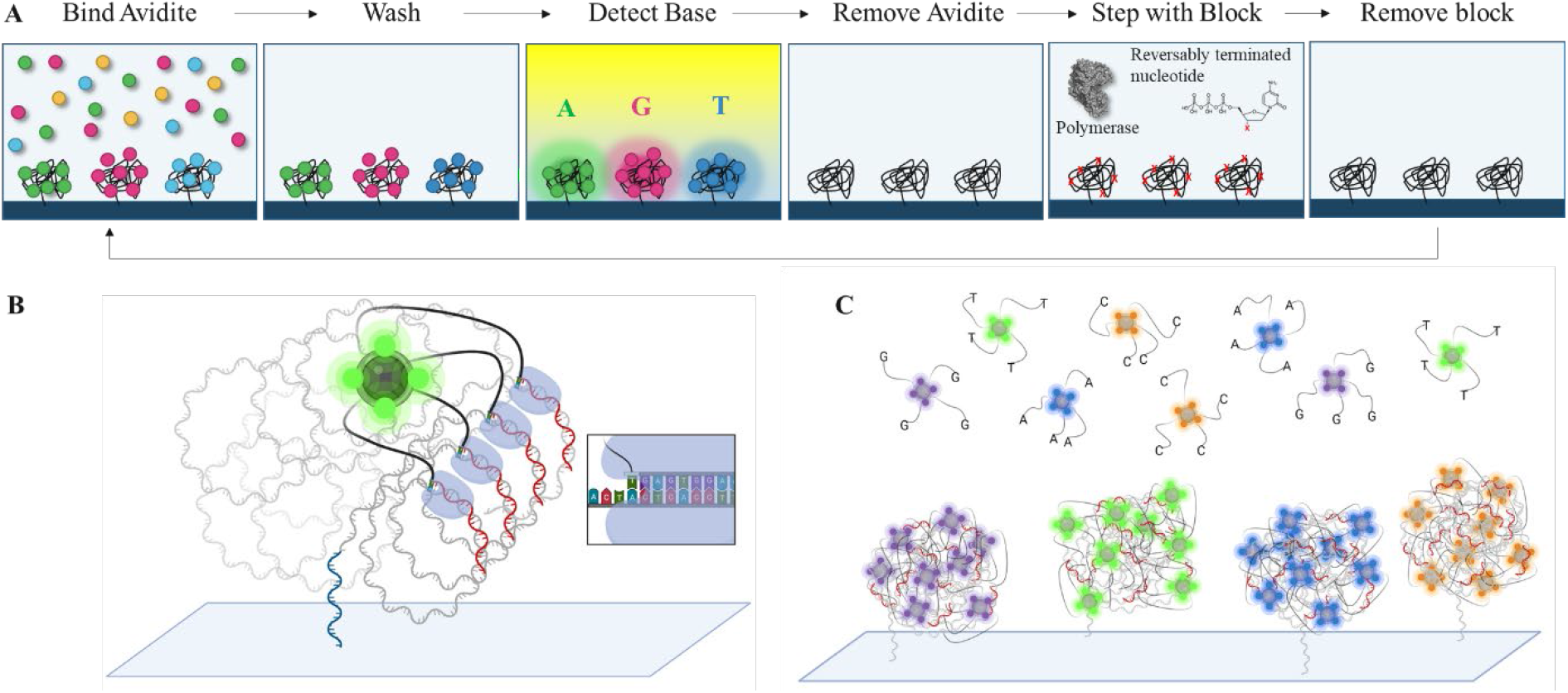
(A) Sequencing by Avidity. A reagent containing multivalent avidite substrates and an engineered polymerase are reacted with DNA polonies inside a flowcell. The engineered polymerase binds to the free 3’ ends of the primer-template of a polony and selects the correct cognate avidite via base-pairing discrimination. The multivalent avidite interacts with multiple polymerases on one polony to create avidity binding that reduces the effective Kd of the avidite substrates 100x compared to a monovalent dye labeled nucleotide, allowing nanomolar concentrations to productively bind. Multiple polymerase-mediated binding events per avidite ensure a long signal persistence time. Imaging of fluorescent, bound avidites enables base resolution. Following detection, avidites are removed from the polonies. Extension by one base using an engineered polymerase incorporates an unlabeled, blocked nucleotide. A terminal 3’ hydroxyl is regenerated on the DNA strand, allowing the cycle to repeat. (B) Rendering of a single avidite bound to a DNA polony via polymerase mediated selection. The initial surface primer used for library hybridization and extension during polony formation is shown in blue. Sequencing primers (red) are shown annealed to the ssDNA polony (grey). Each arm of the avidite (black) connects the avidite core containing multiple fluorophores (green) to a nucleotide substrate. The polymerase bound to the sequencing primer selects the correct nucleotide to base pair with the templating base (inset panel). The result is multiple base-mediated anchor points non-covalently attaching the avidite to the DNA polony. (C) Rendering of multiple DNA polonies with template-specific avidites bound during the binding step of the cycle (polymerase not shown for simplicity). Many avidites bind to each DNA polony generating a fluorescent signal during detection. Multiple long flexible polymer linkers connect the core to the nucleotide substrates.

Avidity sequencing overcomes the kinetic challenges of generating a signal by incorporation of a dye-labeled monovalent nucleotide. In bulk solution, incorporation of a dye-labeled nucleotide is limited by a specificity constant (k_cat_/K_m_) that governs the observed rate of productive nucleotide binding and incorporation[27]. A specificity constant of 0.54±0.22µM^-1^s^-1^ for monovalent dye labeled nucleotides using an engineered polymerase was observed resulting from a maximum rate of incorporation (k_pol_) of 0.86±0.14s^-1^ and an *apparent* Kd (K_d,app_) of 1.6±0.6µM (Fig. 2A). This apparent K_d_ reflects the K_m_ of a kinetic system not in equilibrium rather than the true K_d_ of the nucleotide substrate [28]. To achieve complete product turnover, this high apparent Kd can be overcome by using increased concentrations of fluorescent nucleotide substrate or allowing longer incorporation time for the reaction to complete. Both paths to overcome this substrate limitation have undesirable consequences of high cost or long cycle time. Together, the use of avidity substrates and DNA polonies that contain many copies of substrate DNA in close proximity generate a binding signal, overcomes the limitations of incorporating a monovalent dye labeled nucleotide.

**Figure 2:**
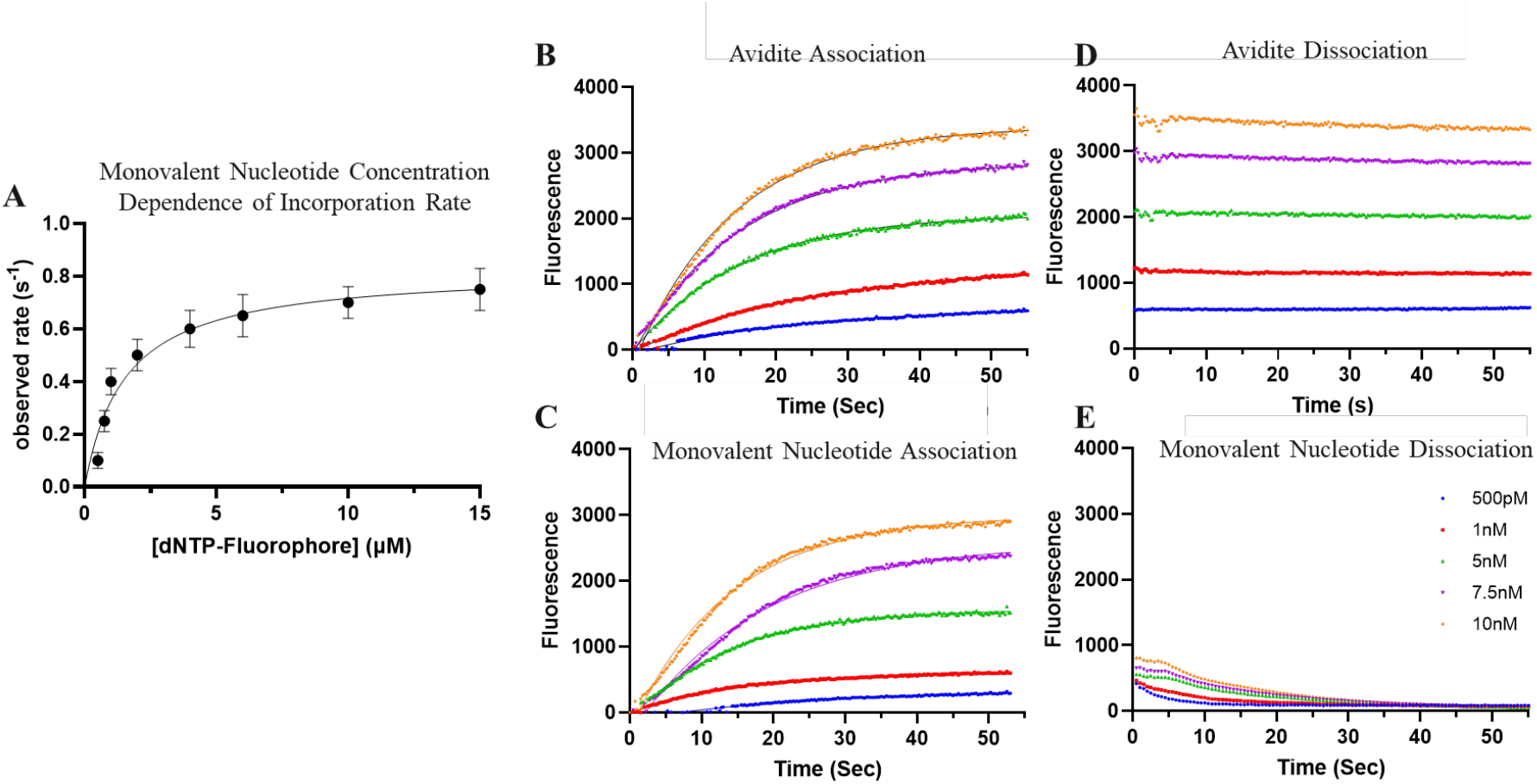
(A) Monovalent fluorophore-labeled nucleotide concentration dependence of the observed rate of incorporation. Time series were performed at each concentration and fit to a single exponential equation to derive a rate. Observed rates were plotted as a function of concentration and fit to a hyperbolic equation to derive a value of k_pol_ = 0.86±0.14s^-1^ and K_d,app_ = 1.6±0.6µM. Real time association kinetics of signal generation resulting from reacting multivalent avidite substrates (B) and monovalent nucleotides (C) with DNA polonies. Real time measurements of signal decay following flow cell washing for imaging of multivalent avidite substrates (D) and monovalent nucleotides (E). Panels B-E use substrate concentrations of 500pM (blue), 1nM (red), 5nM (green), 7.5nM (purple), and 10nM (orange).

Utilizing binding of the four labeled avidites for base identification established a binding equilibrium that reached saturation based on substrate concentration within 30 seconds to generate signal, rather than relying on catalysis. The binding kinetics of this interaction were monitored using real-time data collection to observe avidites binding to polonies with an association rate (k_on,avidite_) of 271±82nM^-1^s^-1^ (Fig 2B). This observed association occurred within the limit of error of a single fluorescently labeled monovalent nucleotide (Fig. 2C). Major differences were observed in the dissociation kinetics of avidite substrates versus monovalent nucleotides. Avidite substrates bound to the DNA polonies tightly with no measurable dissociation over the > 1 minute timescale needed for imaging and basecalling (Fig. 2D). This is in sharp contrast to fluorescently labeled monovalent nucleotides, which dissociated rapidly during the wash step following binding and then continued to dissociate during imaging (Fig. 2E). The negligible dissociation rate resulted in decreased Kd of more than two orders of magnitude for avidites compared to monovalent nucleotides. With the near-zero avidite dissociation-rates, a persistent signal was achieved without the presence of free avidites in bulk solution, eliminating background. Without avidity, dissociation kinetics with monovalent nucleotides showed a 4x signal decrease at the beginning of imaging due to fast dissociation as a result of the disruption of the binding equilibrium during reagent exchange (Fig. 4E).

### Sequencing Instrumentation

Avidity sequencing was performed on the AVITI™ commercial sequencing system. Briefly, the instrument is a 4-color optical system that has 2 excitation lines of approximately 532 nm and 635 nm. The 4-color system is made by using an objective lens, multiple tube lenses and multiple cameras to simultaneously image 4 spectrally separated colors. The detection channels for emission are centered at approximately 532, 570, 635, and 670 nm, respectively. Reagents are delivered using a selector valve and syringe pump to perform reagent cycling. The instrument contains two fluidics modules and a shared imaging module, enabling two flowcells to be utilized in parallel. Subsequent to image collection, data is streamed through an onboard processing unit that performs image registration, intensity extraction and correction, base calling, and quality score assignment as described in the methods section.

### Accuracy of Avidity Sequencing

To evaluate the accuracy of avidity sequencing, 20 sequencing runs were performed using a well characterized human genome. The sequencing data was used to train quality tables according to the methods of Ewing et al. [29], but with modified predictors. The quality tables were then applied to independent sequencing runs. Figure 3 shows the data quality that was obtained in a representative run not used for training. The quality scores were well calibrated across the entire range, meaning that predicted quality matched observed quality as determined by alignment to a known reference. Combined over read 1 and read 2, 96.2% of base calls were above Q30 (an average of one error per 1000 bp) and 85.4% of base calls were above Q40 (an average of one error per 10,000 bp), with a maximum of Q44. For comparison, a publicly available PCR-free NextSeq 500 data set was downloaded from the short read archive[30]. Supplemental figure 1 shows the predicted and observed quality scores. The predicted scores accurately reflected the recalibrated scores. 85.7% of base calls exceeded Q30 and none of the base calls exceeded Q40.

**Figure 3:**
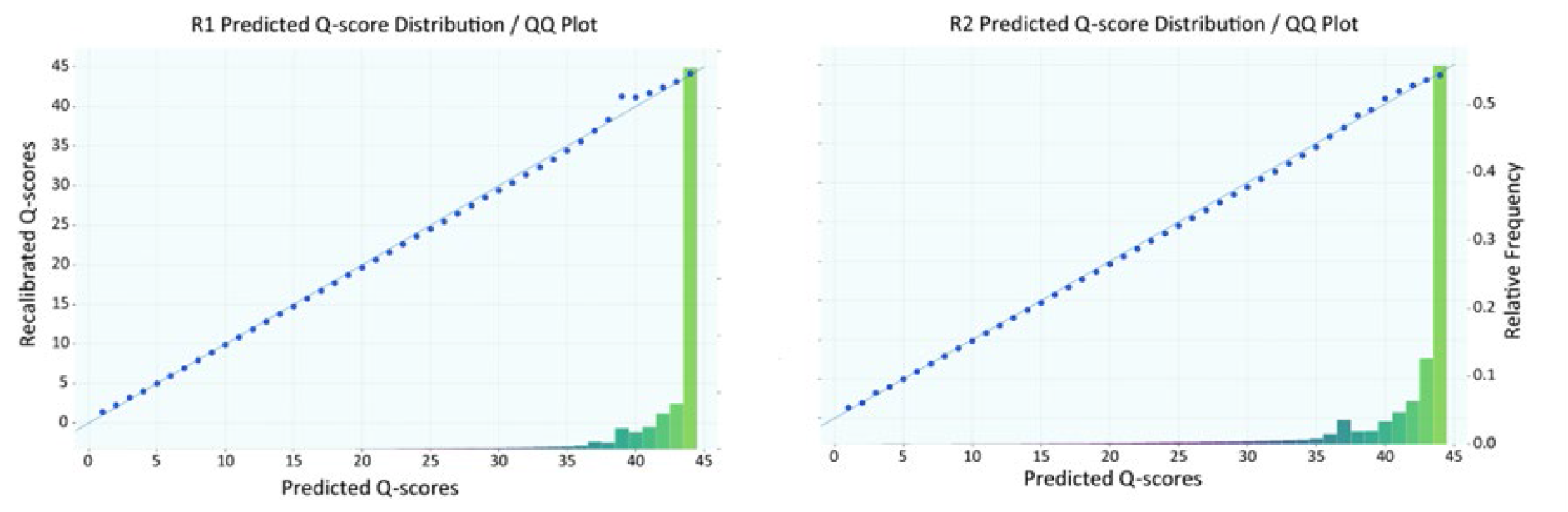
Predicted and observed quality scores for a 2×150 bp sequencing run of human genome HG002. The left panel shows read 1 and the right panel shows read 2. Points on the diagonal indicate that predicted scores match observed scores. The histograms show that the majority of the data points are above Q40, or 1 error in 10,000 bp.

**Figure 4:**
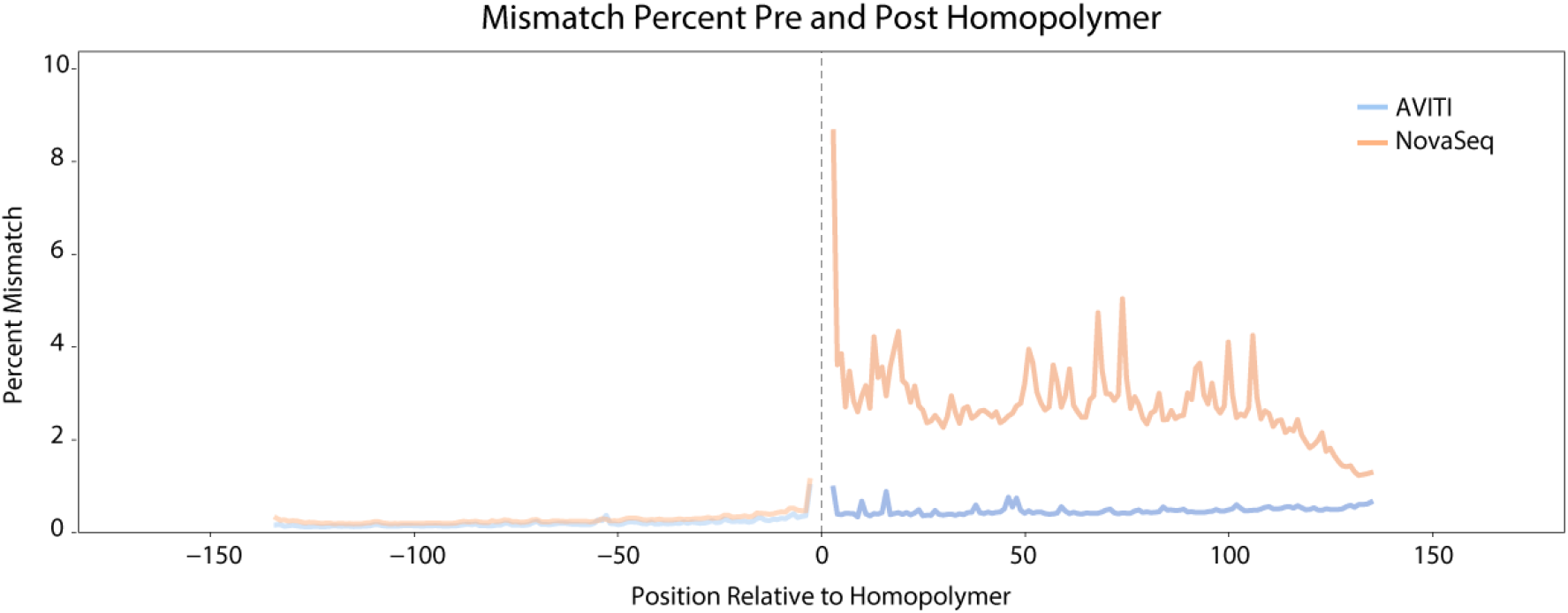
The mismatch percentage of AVITI and NovaSeq reads before and after homopolymers of length 12 or greater.

### Homopolymer sequencing

Sequencing through long homopolymers has posed challenges for multiple sequencing technologies [31, 32]. Although SBS improves homopolymer sequencing relative to flow-based technologies, the error rates of reads that pass through long homopolymer regions increase significantly [33]. Correction algorithms have been proposed to circumvent the inherent challenges with base-calling post-homopolymer repeats[34], but the exact cause has not been fully established in the literature. In contrast to SBS, avidity sequencing leverages rolling circle amplification, polymerases evolved to accommodate the avidite complex formation, and a separate polymerase evolved to efficiently incorporate unlabeled and 3’ blocked nucleotides. We evaluated the impact of these differences on sequencing through long homopolymers. Specifically, homopolymers of length 12 or more nucleotides were used to assess the accuracy of reads before and after homopolymer regions. Figure 4 shows the results comparing avidity sequencing to SBS, averaged across the ~ 700,000 homopolymer loci of length 12 or more. Supplementary figure 1 shows additional runs and additional instruments. Average error rate of avidity sequencing remained stable following a long homopolymer (controlling for the fact that the post-homopolymer stretch occurs in later cycles of a read). By contrast, the error rate of SBS reads increased by more than a factor of 5 following the homopolymer stretches. Figure 5 shows the histogram of pairwise error rate differences between avidity sequencing and SBS for all long homopolymer loci. The avidity sequencing error rate is lower for 97% of the cases and the magnitude of difference is correlated with the homopolymer length. Supplementary figures 2, 3, and 4 show representative loci from the 95^th^, 50^th^, and 5^th^ percentiles of the histogram.

**Figure 5:**
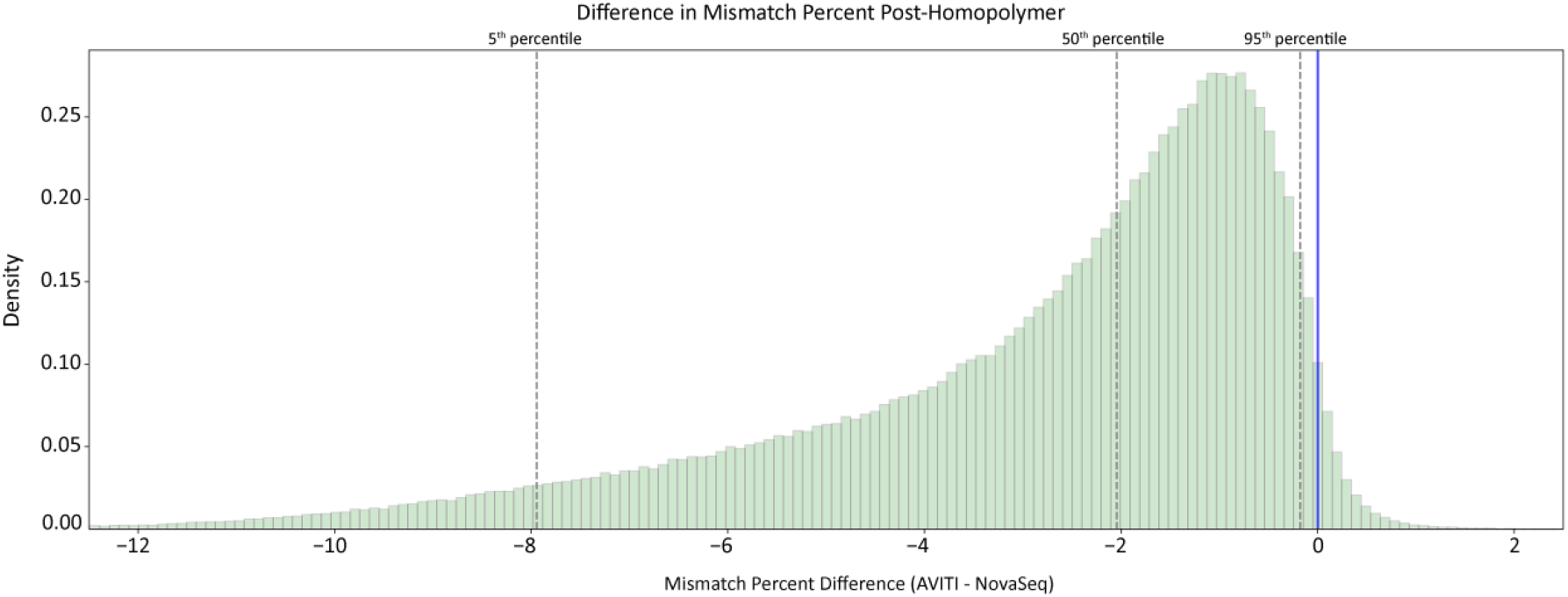
Histogram of pairwise error differences. Difference was selected as the metric to cancel the effects of human variants from the mismatch percent.

### Single Cell RNA Sequencing

To demonstrate sequencing performance across common applications, single cell RNA expression libraries were prepared and sequenced. Two libraries from a reference standard consisting of human peripheral blood mononuclear cells (PBMCs) were generated using the 10X Chromium instrument. The two libraries contain RNA from roughly 10,000 and 1,000 cells, respectively. Following circularization, the libraries were sequenced to generate paired end reads, with read lengths of 28 and 90 for read 1 and read 2, respectively, as recommended by the vendor. The analysis was done using CellRanger [35]. This reference standard is used by 10X Genomics to evaluate sequencing performance, so a set of metrics and guidelines to assess sequencing results is provided along with the biological material. Table 1 shows each metric, the guideline values from 10X Genomics, and the performance of each sequenced library. All metrics were within the guided ranges, and the metrics pertaining to sequencing quality exceeded the thresholds provided.

**Table 1:** Single cell expression: CellRanger metric values for 10K cell and 1K cell libraries from the PBMC reference

| CellRanger v7.0 Metric | Performance expectation | AVITI 10K cells | AVITI 1K cells |
| --- | --- | --- | --- |
| Valid barcodes | >90% | 97.5% | 97.5% |
| Reads mapped confidently to exonic regions | >50% | 53.0% | 53.8% |
| Read mapped confidently to transcriptome | >40% | 74.7% | 77.8% |
| Fraction reads in cells | >80% | 95.5% | 92.6% |
| Q30 bases in barcode | >85% | 99.5% | 99.5% |
| Q30 bases in RNA read | >75% | 98.6% | 98.8% |
| Mean reads per cell | >50,000 | 61,326 | 68,766 |
| Median genes per cell | >1700 | 2,910 | 2,951 |
| Estimated number of cells | +/-20% | 8,513 | 922 |

### Whole Human Genome Sequencing

Another common application is human whole genome sequencing. This application challenges sequencer accuracy to a greater extent than measuring gene expression because the latter requires only accurate alignment while the former depends on nucleotide accuracy to resolve variant calls. To demonstrate performance for this application, the well characterized human sample HG002 was prepared for sequencing using a Covaris shearing and PCR-free library preparation method and sequenced with 2×150bp reads. The run generated 1.02 billion passing filter paired-end reads with a duplicate rate of 0.58% (0.11% classified as optical duplicates by Picard[36]).

A FASTQ file with the base calls and quality scores was down-sampled to 35X coverage and used as an input into the DNAScope analysis pipeline from Sentieon. SNP and indel calls achieved F1 scores of 0.995 and 0.996, respectively. Table 2 shows variant calling performance for SNPs and small indels on the GIAB-HC regions. Sensitivity, precision, and F1-score are shown. The performance on SNPs and indels is comparable.

**Table 2:**
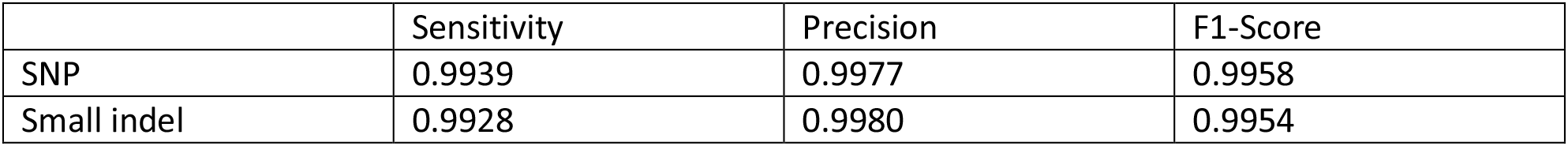
Variant calling performance for HG002 on GIAB-HC regions

### Extensibility of avidity sequencing

Because avidity sequencing is new, many improvement directions have yet to be explored. To assess the extensibility of the chemistry we continued a sequencing run beyond 150 bp to generate a 1×300 dataset from an *E. coli* library. To achieve this, we used a newly developed polymerase and optimized reagent formulation. Figure 6A shows the quality scores as a function of sequencing cycle. Because the quality scores were not trained to these lengths, the scores are approximate. Figure 6B shows the *E. coli* error rate as a function of cycle number based on alignment to the known reference strain. The error rate of the last cycle was 1.9% and the error rate at cycle 150 was 0.1%. Notably, the error calculations were based on the vast majority of the data with a pass filter rate for the run of over 99.6% and BWA settings aimed to strongly discourage soft clipping (no cycles with soft clipping above 0.04%). The enzymes and formulations developed for this run will be leveraged as we continue to identify extensions and improvements.

**Figure 6:**
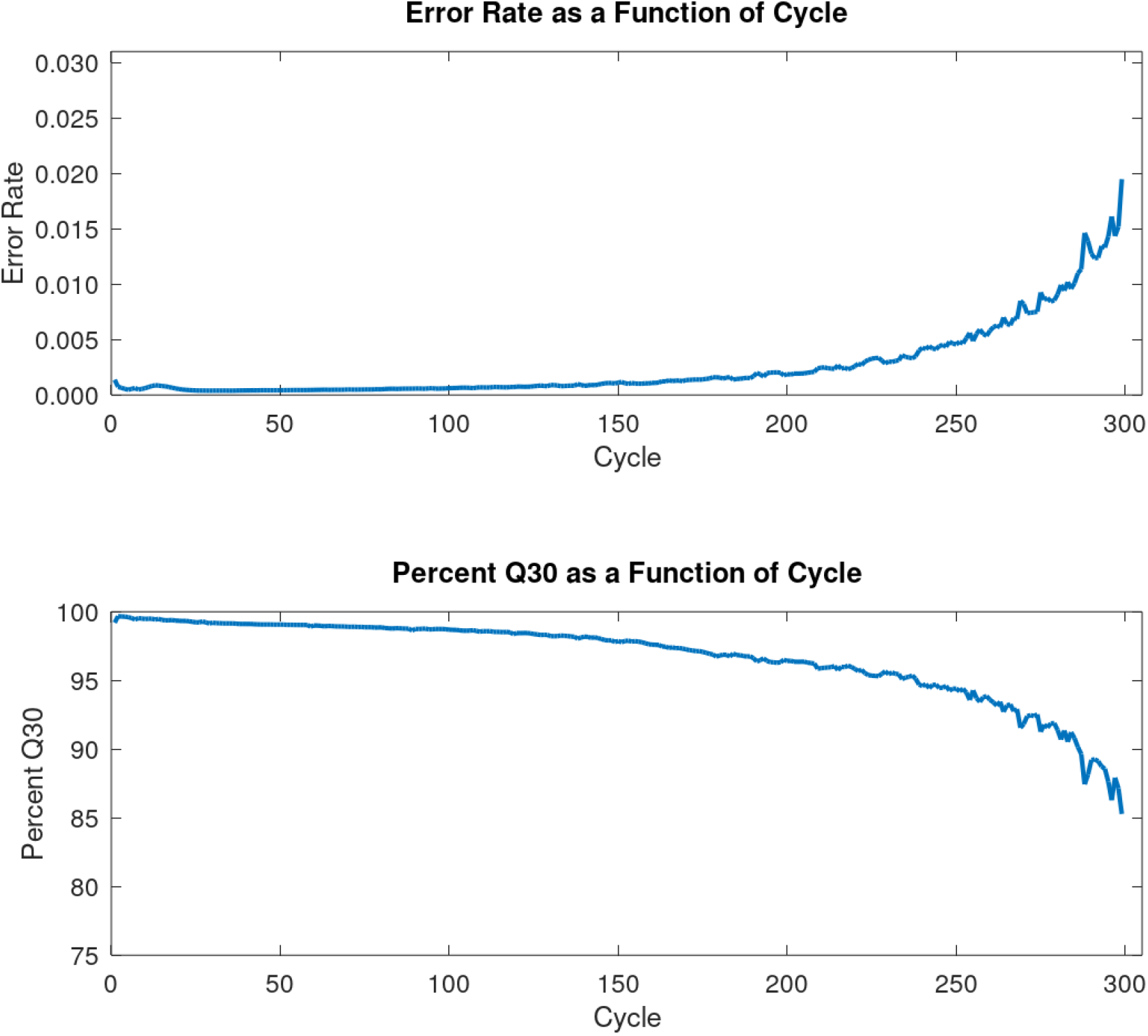
(A) percent Q30 by cycle. Overall Q30 percentage exceeds 96% and end of read has 85% Q30. (B) The E. coli error rate as a function of cycle. Alignment settings strongly discourage soft clipping and greater than 99% of reads pass filter. Last cycle error rate was 0.019.

## Discussion

We have presented a new sequencing chemistry that achieves improved quality and lower reagent consumption by independently optimizing nucleotide incorporation and signal generation. Although other chemistries have proposed to separate incorporation and signal generation [37], the avidite concept is new and benefits from the fact that multiple nucleotides on the avidite bind multiple copies of the DNA template within a polony. Furthermore, the avidite construct is modular. The core can be swapped for a different substrate. The number and type of dye molecules are configurable, and many types of linkers can be used. The changes are straightforward to implement and do not require modification to the polymerase responsible for binding the nucleotides attached to the linkers. The modular design speeds technology improvement as each component can be optimized in parallel for increased signal, decreased cycle time, lower reagent concentration, or any other potential axis of improvement.

The avidity chemistry described above has been implemented as part of a benchtop sequencing solution. The accuracy of the sequencer has been demonstrated by training a quality model on human sequencing data, which shows that that majority of bases in an independent human whole genome sequencing run exceed Q40, or less than 1 error in 10,000 base pairs. The high accuracy likely results from the use of an engineered high-fidelity polymerase, synergistic binding of multiple nucleotides on a single avidite to ensure only the correct cognate avidite binds to the polony, and a binding disadvantage for out-of-phase DNA copies inside of a polony that lack other out-of-phase neighbors to serve as avidity substrates. In addition to overall accuracy improvements, the chemistry retains good performance in reads that contain long homopolymers. The sequencer can be used on a wide range of applications, as exemplified by results for single cell RNA-seq and for whole human genome sequencing. In both cases, reference standards were sequenced so that the quality of result could be assessed. The single-cell data exceeded the quality metric guidelines provided by 10X Genomics[38]. The human genome variant calling results showed high sensitivity and precision for both SNPs and small indels [39]. The two applications were selected due to the availability of well-characterized samples and because they represent very different use cases. However, these are only examples and many other applications have already been demonstrated, with manuscripts in preparation for targeted sequencing, metagenomics, low pass sequencing, and methylation sequencing applications[40]. Notably, the current implementation of the avidity-based chemistry is new. Although it already achieves high accuracy and broad applicability, there are many improvement directions being explored. In addition to the initial demonstration of longer reads shown here, further quality improvements, shorter cycle times, and higher densities are under development.

## Methods

### Solution measurements of nucleotide incorporation

Solution measurements of nucleotide kinetics were performed using commercially available dGTP-Cy5. DNA substrates for solution kinetic assays were prepared by annealing a 5’FAM labeled primer oligo purchased from IDT and HPLC purified (5’GCAGCCGTCCATCCTACTCA3’) with a template oligo (5’ACGACCATGATGAGGATGGACGGCTGC3’). Annealing was performed with 10 percent excess template oligo in annealing buffer using a PCR machine to heat oligos to 95°C followed by slow cooling to room temperature over 60 minutes. Solution kinetics were performed by mixing a preformed Enzyme-DNA complex with fluorescent nucleotide and MgSO_4_ using a RQF3 Rapid Quench Flow (KinTek Corp.). The enzyme used was an engineered variant of Candidatus altiarchaeales archaeon. The final reaction was conducted in 25mM Tris pH 8.5, 40mM NaCl, 10mM ammonium chloride at 37C. Extension products were separated from unextended primer oligos by capillary electrophoresis using a 3500 Series Genetic Analyzer (ThermoFisher) to achieve single base resolution. Products were quantified and fit to a single exponential equation. The observed rates as a function of nucleotide concentration were then fit to a hyperbolic equation to derive an *apparent* K_d_ (K_d,app_) and a rate of polymerization (k_pol_).

### Avidite synthesis and construction

Initial research scale avidites were constructed by dissolving 5mg 10kD 4-arm-PEG-SG (Laysan Bio Item# 4arm-PEG-SG-10K-5g) into 100 uL 95% organic solvent (e.g., ethanol) 5 mM MOPS pH 8.0 to make a 50 mg/mL solution (5 mM). 19uL of the solution was combined with 1.5uL 10mM dATP-NH_2_ (7-Deaza-7-Propargylamino-2’-deoxyadenosine-5’-Triphosphate from Trilink N-2068), 8.0uL 3.75mM 2kD Biotin-PEG-NH_2_ (Laysan Bio Item# Biotin-PEG-NH2-2K-1g) in 95% organic solvent (e.g., ethanol) 5 mM MOPS pH 8.0. After mixing, 5mM 10kD 4-arm-PEG-SG was added. The final composition was 0.50mM dA-NH_2_, 1.0mM biotin-PEG-NH2 (2kD), 0.25mM 4-arm-PEG-NHS, 85.5% organic solvent (e.g., ethanol), 4.5mM MOPS pH 8.0. Following a 1000rpm incubation at 25C for 90 minutes, the reaction volume was adjusted to 100uL by addition of MOPS pH 8.0. Purification was performed using a Biorad Biospin P6 column pre-equilibrated in 10mM MOPS pH 8.0. The purified dATP-PEG-Biotin complex was mixed with Zymax Cy5 Streptavidin in a 2.5:1 volumetric ratio and allowed to equilibrate for 30 minutes at room temperature.

### Real-time measurements of avidite association and dissociation

Real-time measurement of avidite binding kinetics were performed using an Olympus IX83 microscope equipped with automated fluidics, excitation bands of 532 and 635 nm, dichroics and filter detection apparatus for emission detection at peak wavelengths of 532, 570, 635 and 670 nm, respectively, and custom control software. Flowrates of 60uL/s were used for reagent exchanges. The instrument and flowcells were used to amplify genomic DNA. The instrument was paused following polony generation and priming and the flowcell was moved to the microscope. Data collection (4 fps) was triggered by flow of the avidity mix and collected for 55 seconds. Polonies in the field were localized by spot finding algorithm and background corrected intensities were extracted vs time. Experiments were performed at 0.5 pM, 1 nM, 7.5 nM, and 10 nM avidite or monovalent dye labeled nucleotide concentrations. Substrates at the respective concentrations were combined with 100 nM of the engineered enzyme variant of Candidatus altiarchaeales archaeon in the commercially available avidity buffer formulation. Avidites or nucleotides were labeled with Alexa Fluor 647. Higher concentration data collection was limited by the ability to detect polony intensity from free avidite intensity at elevated concentrations. Off-rate measurements were performed by binding avidites to flowcell polonies, followed by washing with imaging mix and triggering data collection.

### Genomic DNA and NGS library preparation

Human DNA from cell line sample HG002 was obtained from Coriell Institute. Linear NGS library construction was performed using a KAPA HyperPrep library kit (Roche) according to published protocols. Finished linear libraries were circularized using Element Adept Compatibility kit (catalog #830-00003). Final circular libraries were quantified by qPCR with the standard and primer set provided in the kit. Circular library DNA was denatured using sodium hydroxide and neutralized with excess Tris pH 7.0 prior to dilution. Denatured libraries were diluted to 8pM in hybridization buffer before loading onto the sequencing cartridge.

### Single cell 3’ gene expression library circularization

Single cell RNA-Seq libraries were prepared from two lots of PBMC cell suspensions (10,000 cells and 1,000 cells) using the Chromium Next GEM Single Cell 3’ Kit v3.1 (Part #1000268). Each library was quantified and individually processed for sequencing using the Adept Library Compatibility Kit (Part #830-00003). The processed libraries were pooled and sequenced with 28 cycles for Read 1, 90 cycles for Read 2, and index reads.

### Sequencing instrument and workflow

Element’s AVITI commercial system (Part #88-00001) was used for all sequencing data. AVITI 2×150 kits were loaded on the instrument (Part #86-00001). Primary analysis was performed onboard the AVITI sequencing instrument and FASTQ files were subsequently analyzed using a secondary analysis pipeline from Sentieon.

### Sequencing primary analysis

Four images were generated per field of view during each sequencing cycle, corresponding to the dyes used to label each avidite. An analysis pipeline was developed that uses the images as input to identify the polonies present on the flowcell and to assign to each polony a base call and a quality score for each cycle, representing the accuracy of the underlying call. The analysis approach has similar steps to those described in Whiteford et al. [25]. Briefly, intensity is extracted for each polony in each color channel. The intensities are corrected for color cross talk and phasing. The intensities are then normalized to make cross channel comparisons. The highest normalized intensity value for each polony in each cycle determines the base call. In addition to assigning a basecall, a quality score corresponding to the call confidences is also assigned. The standard Q score definition is utilized, where the Q value is defined as *Q* = − 10 * log_10_ *pp*, where p is the probability that the base call is an error. The Q score generation follows the approach of Ewing et al., with modified predictors [21], and is encoded using the phred+33 ASCII scheme. The sequence of base call assignments and quality scores across the cycles constitute the output of the run. This data is represented in standard FASTQ format for compatibility with downstream tools.

### Quality score assessment

To assess the accuracy of quality scores shown in Figure 3, the FASTQ files were aligned with BWA to generate BAM files. GATK BaseRecalibrartor was then applied to the BAM, specifying publicly available known sites files to exclude human variant positions.

### Homopolymer analysis

A BED file provided by NIST genome-stratifications v3.0, containing 673,650 homopolymers of length greater than 11 was used to define the regions of interest for the homopolymer analysis (GRCh38_SimpleRepeat_homopolymer_gt11_slop5). Reads that overlapped these BED intervals (using samtools view -L and adjusting for the slop5) were selected for accuracy analysis. Reads with any of the following flags set were discarded (secondary, supplementary, unmapped or reads with mapping quality of 0). Reads were oriented in the 5’ -> 3’ direction, and split into 3 segments, preceding the homopolymer, overlapping the homopolymer, and following the homopolymer. The mismatch rate for each read-segment was computed, excluding N-calls, softclipped bases and indels. For example, if a 150 bp read (aligned on the forward strand) contains a homopolymer in positions 100-120, then the first 99 cycles were used to compute the error rate prior to the homopolymer, and the last 30 cycles were used to compute the error rate following the homopolymer. Reads were discarded if either the sequence preceding or following the homopolymer was less than 5bp in length. All reads were then stacked into a matrix, according to their positional offset relative to the homopolymer, and error rate per pos-offset was computed.

The average error rate was computed for avidity sequencing runs and for publicly available data from multiple SBS instruments, for comparison. The differences of mismatch percentages, across all BED intervals, between AVITI™ and NovaSeq were plotted in a histogram and examples showing various percentiles within the distribution were chosen for display via IGV.

### Single cell gene expression data analysis

Following sequencing, the Bases2Fastq Software was used to generate FASTQ files for compatible upload into 10x Cloud and subsequent analysis with the 10x Genomics Cell Ranger analysis package. Data visualization of single cell gene expression profiling was generated using 10x Genomics Loupe Browser.

### Whole genome sequencing analysis

A FASTQ file with the base calls and quality scores was down-sampled to 35X raw coverage (360,320,126 Input reads) and used as an input into Sentieon BWA following by Sentieon DNAscope [41]. Following alignment and variant calling, the variant calls were compared to the NIST genome in a bottle truth set v4.2.1 via the hap.py comparison framework to derive total error counts and F1 scores[42].

### 1×300 Data generation

An E. coli library was prepared using enzymatic shearing and PCR amplification. The library was then sequenced for 300 cycles using new enzymes for stepping along the DNA template and for avidite binding. The reagent formulation using increased enzyme and nucleotide concentration during the stepping process was used to improve stepping performance. The contact time for avidite binding and the exposure time were both reduced without performance losses to decrease cycle time over the 600 cycles of sequencing. The displays show 299 cycles of data as cycle 300 is only used for prephasing correction. To minimize soft clipping during alignment, the following inputs were used in the call to BWA-MEM: -E 6,6 -L 1000000 -S.

## Data availability

The avidity sequencing data sets described in the manuscript are available for download via the AWS CLI in the following public bucket: s3://avidity-manuscript-data/, pending upload to the short read archive.

## Acknowledgements

We would like to thank Dr. Joseph Puglisi and Dr. Tuval Ben-Yehezkel for valuable comments and discussion during the writing of the manuscript.

