## Supplemental Figures for "Sequencing by avidity enables high accuracy with low reagent consumption"

Supplementary Figures

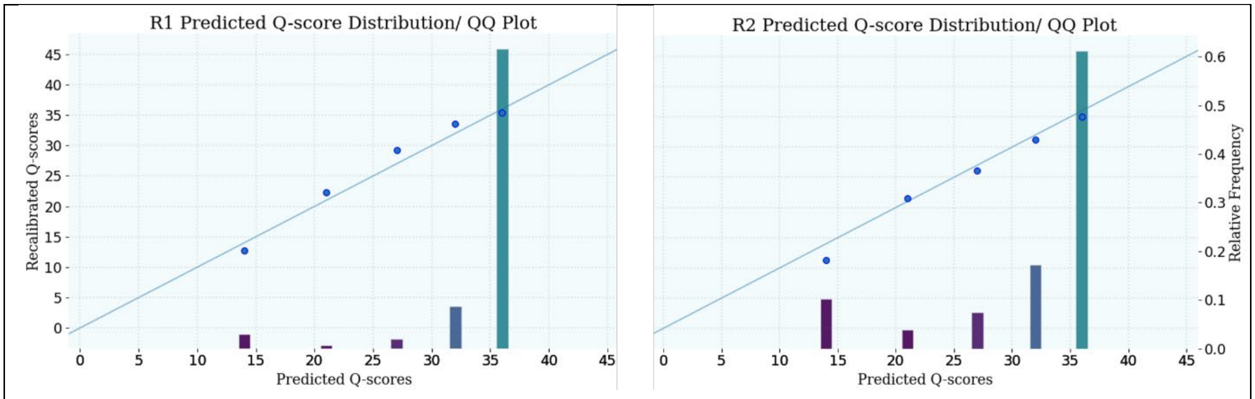


*Supplementary Figure 1: Quality scores for a publicly available 2x150 NextSeq 500 run. The sequenced library is PCR-free HG002. Quality scores are binned to 5 possible values, presumably to maximize data compression.*

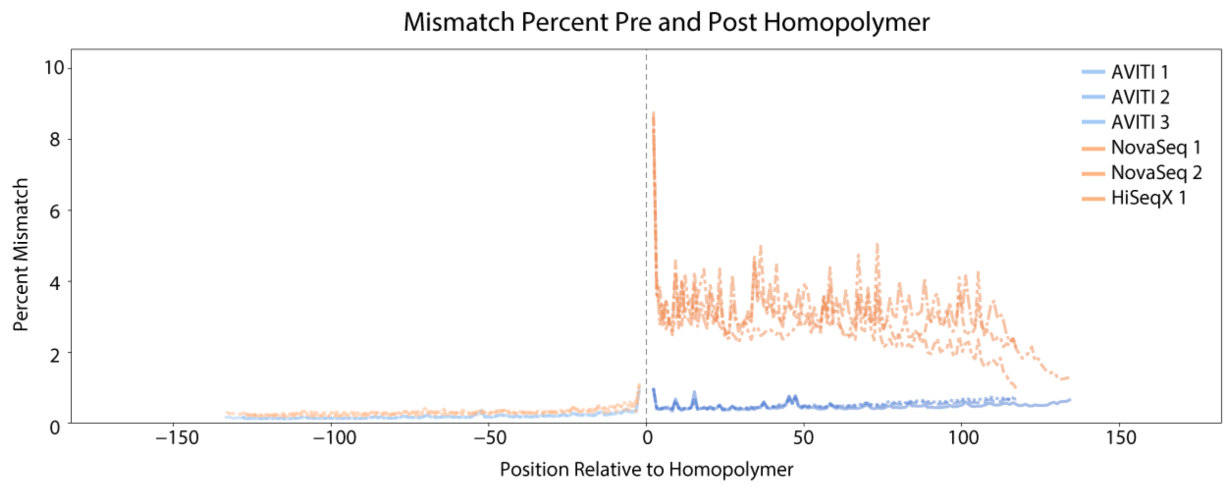


*Supplementary Figure 2: Pre and post homopolymer mismatch percentages across multiple runs and sequencing instruments*

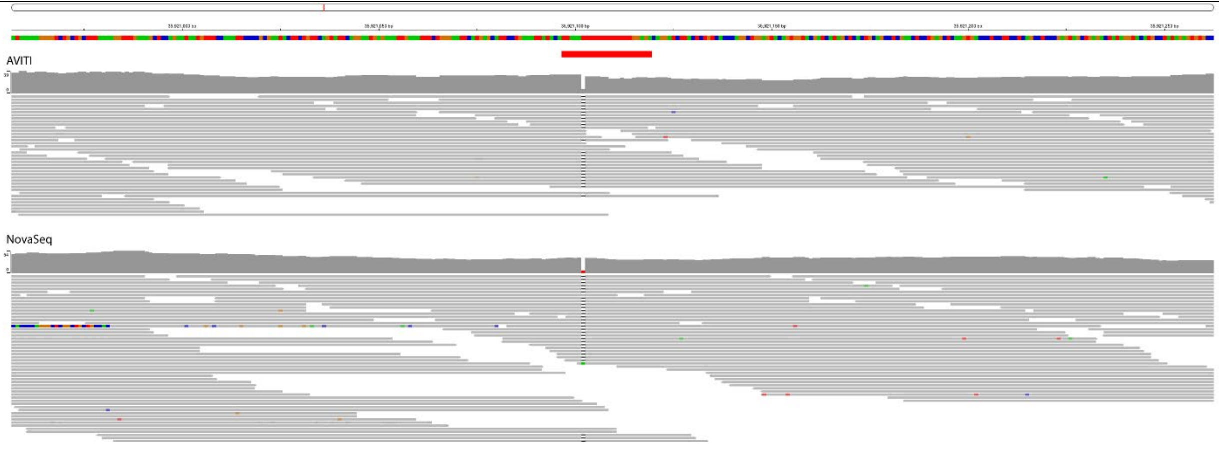


*Supplementary Figure 3: IGV display of a representative homopolymer locus at the 95$^{th}$ percentile of AVITI minus NovaSeq mismatch percent, where AVITI and NovaSeq mismatch percentages are most similar (corresponding to the right-most dashed line of figure 5). The red bar at the top indicates the homopolymer.*

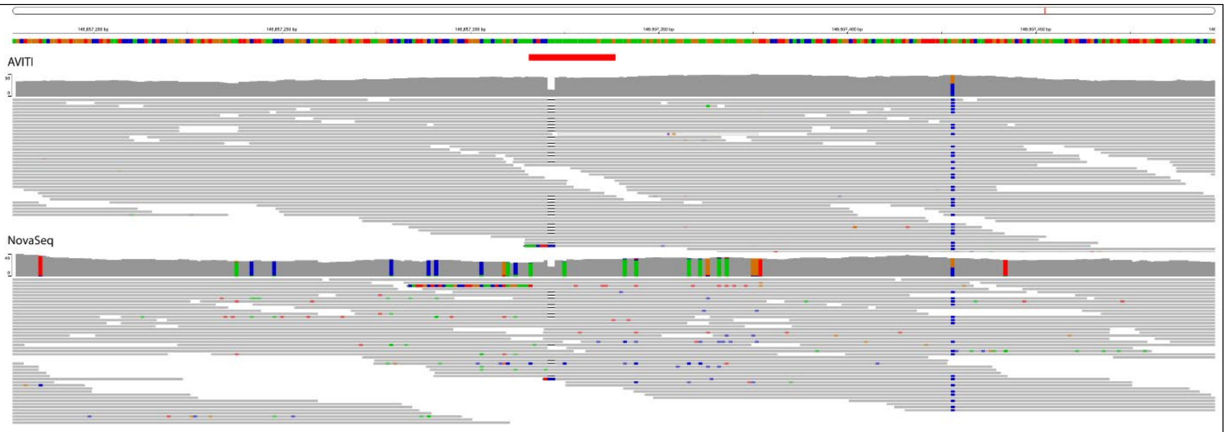


*Supplementary Figure 4: IGV display of a representative homopolymer locus at the 50th$^{th}$ percentile of AVITI minus NovaSeq mismatch percent (corresponding to the middle dashed line of figure 5). The red bar at the top indicates the homopolymer.*

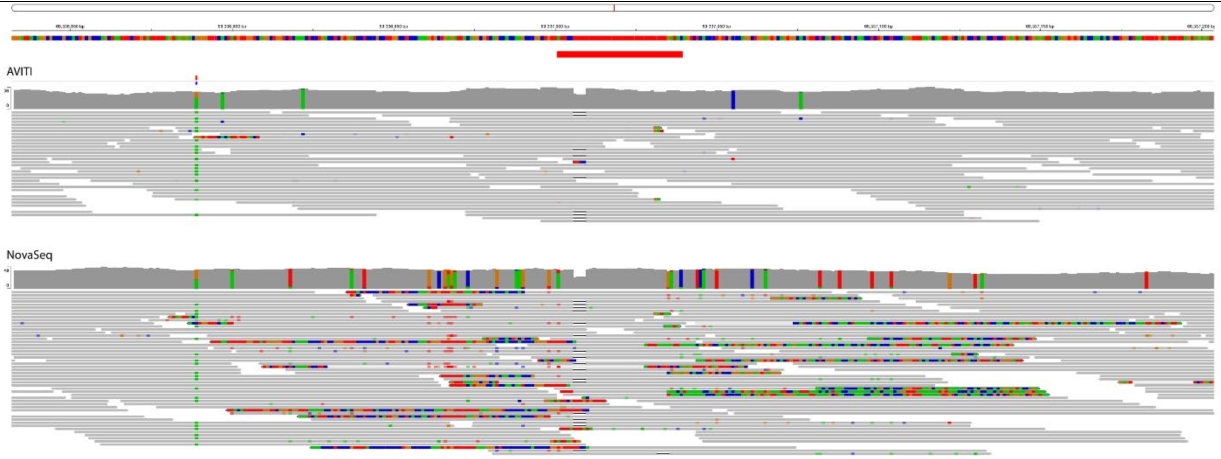


*Supplementary Figure 5: IGV display of a representative homopolymer locus at the 5th$^{th}$ percentile of AVITI minus NovaSeq mismatch percent (corresponding to the left-most dashed line of figure 5). The red bar at the top indicates the homopolymer.*
